# A Computational Pipeline for Retinal Capillary Blood Flow Measurement using Adaptive Optics Line Confocal Ophthalmoscopy

**DOI:** 10.64898/2026.09.21.753337

**Authors:** Syed Muhammad Hamza Shah, Lin Tong, Ruixue Liu, Yuhua Zhang, Jianhua Wang, Liang Liang

## Abstract

**Background and Objective:** High-speed and high-resolution retinal imaging using adaptive optics line confocal ophthalmoscopy (AOLCO) resolves individual red blood cells (erythrocytes) flowing through the smallest retinal capillaries, but quantifying their velocity first requires removing linear and nonlinear distortions caused by eye motion that may result in a displacement of tens to hundreds of pixels between frames. We present a computational pipeline that stabilizes AOLCO capillary video and measures erythrocyte velocity in true physical units, delivered as an interactive 3D Slicer extension and a batch command-line tool sharing one analysis core.

**Methods:** Our video analysis pipeline has four stages: preprocessing, registration, postprocessing, and velocity measurement. During preprocessing, illumination is corrected, and closed-eye frames are identified. During registration, each frame is aligned to a reference frame using affine and, when needed, B-Spline registration, initialized by our robust and fast (GPU-based) translation-registration algorithm to accommodate large eye movements (e.g., microsaccades). During postprocessing, adjacent registered frames are differenced to suppress stationary structures, and a 2D projection of the frame-difference video reveals the capillary network. During velocity measurement, an operator manually identifies capillary segments on the 2D projection image. Spatiotemporal images are then generated along these segments from the frame-difference video, and blood-flow velocity curves are obtained from these images using the Radon transform.

**Results:** We validate the velocimetry of the pipeline against synthetic ground-truth flow data, and the experiments show that the velocimetry recovers the speed within an error of ±4% The GPU-based translation registration algorithm aligns the large eye motions (∼90 µm per frame on average) and runs about 3.6× (100 Hz) to 38× (400 Hz) faster than the baseline method. Across the full cohort (2344 videos, 57 subjects, 30 Hz to 400 Hz), our registration approach improves structural similarity (SSIM) of the video frames on every acquisition from 0.73 → 0.86. The recovered blood flow velocities are physiologically plausible, of the same order as published AO measurements.

**Conclusions:** Our computational pipeline enables AOLCO velocimetry on the capillary-level.

## 1. Introduction

The retinal microcirculation feeds the high metabolic demand of the inner retina, and its breakdown is an early sign of diabetic retinopathy and is linked to glaucoma, age-related macular degeneration, and systemic conditions such as hypertension, stroke, and Alzheimer’s disease [1]. Measuring blood flow in single capillaries, where erythrocytes squeeze through lumens only ∼5 µm wide, would give a direct, non-invasive window onto this physiology. Most established techniques — laser Doppler velocimetry, optical coherence tomography angiography (OCTA) [2], laser speckle imaging — map vessel structure or bulk perfusion well, but lack either the spatial resolution to isolate one capillary or the temporal resolution to follow the cells inside it.

Adaptive optics (AO) ophthalmoscopy cancels the eye’s own optical blur and resolves the smallest capillaries [3]. At a high frame rate it can image moving erythrocytes and, in principle, recover their velocity from the slope of the streaks they trace in a fixed vessel [1]. Two obstacles stand between a raw video and a velocity number. First, the eye is never still: tremor, drift, micro-saccades, and blinks move the retinal image by tens to hundreds of pixels from one frame to the next, so the vessel does not stay put and no per-pixel time series can be read until the video is stabilized. Second, even after stabilization, turning the streak slope into a physical speed in µm*/*s couples the camera’s pixel pitch, the true frame rate, and the streak orientation, and a small error in any of these passes straight through to the reported velocity.

This paper presents an open, reproducible pipeline that addresses both obstacles. Its contributions are: (i) a faithful, open *Radon-based velocimetry* stage that reports true µm*/*s and is *validated against synthetic ground truth*, with an explicit pixel-pitch calibration that a pixels-per-frame estimator omits; (ii) a *GPU patch-based translation registration* for high-speed AO video, evaluated by its inter-frame self-consistency and *benchmarked against gradient-based itk-elastix translation* in speed and robustness to large eye motion; (iii) an optional *hierarchical elastix refinement* (translation → affine → B-spline) that composes transforms instead of resampling between stages; and (iv) an *open packaging* of the whole pipeline behind one analysis core, exposed through both a 3D Slicer extension and a batch tool, demonstrated on real acquisitions. Together these make capillary-level velocimetry reproducible at cohort scale — a prerequisite for using retinal hemodynamics as a clinical biomarker.

The novelty is integrative rather than algorithmic, and we state this plainly. The individual primitives are established and cited as such — normalized cross-correlation [4], RANSAC [5], elastix B-spline registration [6, 7], and the flat-field Radon estimator of Duncan et al. [8]. The contribution is their assembly into an open, GPU, whole-frame pipeline with a dual 3D Slicer/batch surface over one analysis core, together with the first reproducible head-to-head of whole-frame patch registration against an open gradient-based baseline (Section 4.3) and an explicit, synthetic-validated pixel-pitch calibration that a pixels-per-frame estimator omits.

## 2. Related work

### Registration

Aligning a moving window to a reference is most often posed as a normalized cross-correlation match [4, 9], with outlier matches rejected by a robust estimator such as RANSAC [5, 10]. A parallel family estimates the same shift in the Fourier domain by phase correlation [11, 12], refined to sub-pixel accuracy either by upsampled cross-correlation [13] or by intensity-based pyramid schemes [14, 15]; these are fast but estimate one global shift per frame and offer no natural outlier rejection when part of the field is occluded, which is the regime that dominates high-speed retinal video. Residual non-rigid distortion is then removed by intensity-based deformable registration, typically B-spline free-form deformations [7] as packaged in elastix [6]. A dedicated line of work targets adaptive optics scanning laser ophthalmoscopy, where strip-based methods register sub-frame strips to correct eye motion [16, 17], and where the same strip correlation can be run in real time as an optical eye tracker [18]. General-purpose registration toolkits — ANTs [19], SimpleITK [20] and SimpleElastix [21] — and learning-based deformable registration [22] solve a more general problem than ours; here the transform is known to be a per-frame translation, and exploiting that is what makes the GPU stage fast enough for thousand-frame acquisitions. Our registration applies the cross-correlation and robust-averaging primitives to *whole-frame* stabilization of high-speed retinal video on the GPU, and offers an open alternative to the separate strip-based software those studies relied on.

### Velocimetry and vessel mapping

Given a stabilized stack and a vessel tracing, the cell velocity equals the slope of the streaks in the space–time image along the vessel. The Radon transform recovers this slope robustly by finding the projection angle of best alignment [23, 24]. For fundus and AO imaging, Dun-can and colleagues [8] added a flat-field normalization and a root-mean-square angle metric that suppress slow illumination drift; this is the formulation we port and validate. The alternative to a window-aggregate slope is to follow individual objects, as particle tracking [25] and particle image velocimetry [26, 27] do in general. In AO retinal video this has largely been done on the space–time image as well: Tam and Roorda extract the trace of a *single* object and take its slope by linear regression, correcting first for the raster-scan and eye-motion distortion a scanning system imposes on that trace [28, 29]; Bedggood and Metha instead visualize the cell stream directly at high frame rate [30]. A per-object trace gives per-cell speeds but needs each object resolved and followed; the Radon slope needs only that the streaks within a window share an orientation, which is what survives at capillary contrast. Separately, mapping *which* vessels carry flow is the job of motion-contrast imaging, e.g. OCTA from temporal decorrelation or speckle variance of the OCT signal [2, 31]; our vessel-network map uses the same idea in the spatial domain.

### Relation to Gu et al. [1]

That study is our reference instrument family, the workflow we mirror, a source of the Radon principle, and our only in-vivo velocity comparator, so we state the delta plainly. We *keep* the motion-contrast network rendering, the Radon angle principle, and the parafoveal near-confocal target. We *replace* their separate, custom motion-correction software with an open, whole-frame GPU patch matching plus RANSAC and automatic reference-frame selection. We *add* an explicit pixel-pitch calibration validated against synthetic ground truth, and we *package* the pipeline behind one core exposed through both a 3D Slicer extension and a batch tool.

## 3. Methods

### 3.1. Imaging device and pipeline overview

The videos were acquired with a high-speed *adaptive optics near-confocal ophthalmoscope* of the family introduced by Gu et al. [1, 3], which corrects the eye’s wave aberration with a wavefront sensor and deformable mirror to resolve single erythrocytes in the smallest capillaries. It images the parafoveal capillary bed of a dilated, fixating eye at a pixel pitch of *p* = 0.67 µm*/*pixel. The registered cohort spans four acquisition rates (30 Hz to 400 Hz, Table 1); two of them carry the analyses in this paper: 100 Hz over the full 2048 × 2046 field, which renders the perfused vessel network, and 400 Hz over a reduced field height, which resolves erythrocyte motion for velocimetry. The remaining rates appear only in the cohort-wide registration evaluation of Section 4.2.

**Table 1:**
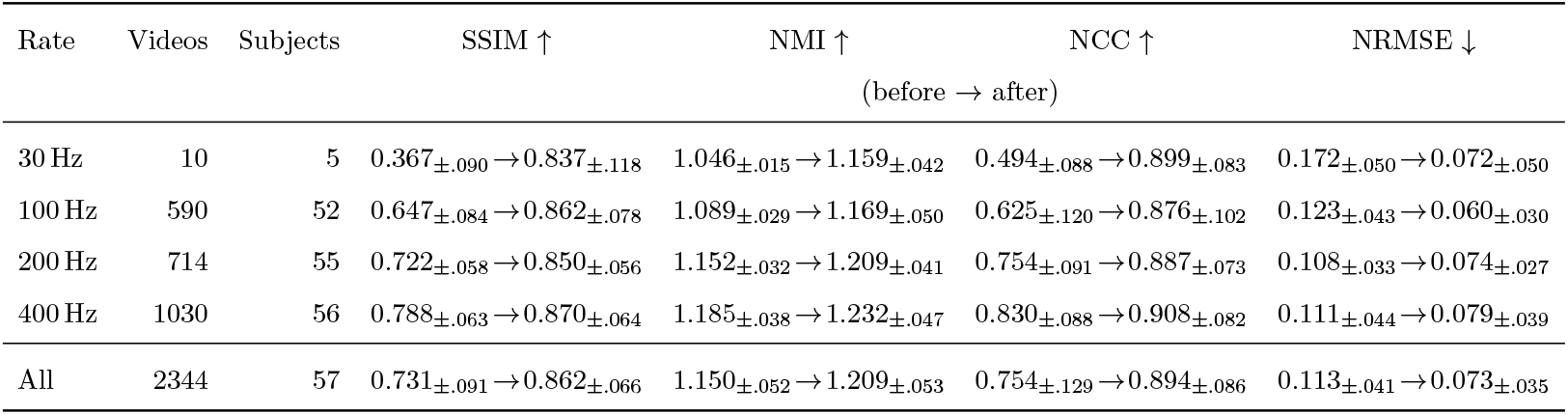
Cohort registration quality (before vs. after), by acquisition rate: mean image similarity over the whole registered cohort (2344 videos from 57 subjects; up to 120 evenly spaced pairs of consecutive eye-open frames per video, central 1024 × 1024 crop clipped to the rows available at 200 and 400 Hz). Cells are the across-video mean, with the across-video *sample* standard deviation as a subscript (NumPy ddof= 1; the distinction is invisible at *n* = 2344 but not in the 30 Hz row, where *n* = 10). “Before” is the re-shift proxy — the registered frames with their recorded inter-frame shift re-injected — so this is an internal self-consistency check, not a comparison against an unregistered baseline or a competing method, and the proxy carries a one-sided interpolation/border penalty. Registration improves every metric (mean) at every rate; SSIM improves on all 2344 videos (minimum gain +0.003) and 2334 improve on all four metrics. Subject counts overlap across rates (many subjects were imaged at several rates); 57 are distinct. Cells pool videos, and subjects contribute unequally (5–91 videos each, median 40); re-weighting so that every subject counts equally moves post-registration SSIM from 0.862 to 0.855 and NCC from 0.894 to 0.888, so the improvement is not carried by the heaviest-contributing subjects.

The instrument is a custom adaptive-optics line-confocal scanning laser ophthalmoscope built in the laboratory [32]. A superluminescent diode (BroadLighter S795-HP, Superlum, Ireland; *λ* = 795 nm, bandwidth 15 nm) is collimated and focused by a cylindrical lens into a line, relayed through the scanning and adaptive optics to form a two-dimensional raster. Backscattered light retraces the incoming path onto a high-speed line camera (OctoPlus EV71Y01, Teledyne DALSA, Canada) whose sensor acts as a one-dimensional confocal gate along the scan direction, with 10 µm × 200 µm pixels. The adaptive optics comprises a custom Shack–Hartmann sensor and two deformable mirrors (DM97-15 and DM192, Bertin Alpao, France) in a woofer–tweeter configuration; aberrations are measured over a 6.75 mm pupil with an 832 nm beacon (SuperK, NKT Photonics, Denmark) at 335 sampling points, and closed-loop correction runs at 50 Hz, typically leaving a residual RMS wave-front error near 0.04 µm (diffraction-limited). The instrument is documented at a 5^◦^ × 5^◦^ retinal field digitized at 2048 × 2048 pixels at its base 30 Hz rate [32], which is the origin of the *p* = 0.67 µm*/*pixel pitch used throughout; the faster read-out modes below keep the same lateral sampling. Imaging and beacon powers are 2.0 mW and 25 µW, a composite exposure of 0.34 of the ANSI maximum permissible limit [32]. The head is stabilized on a chin rest and head mount, with a flashing green target guiding fixation. The two rates used here trade field height for speed: 100 Hz covers the full 5^◦^ × 5^◦^ field at 2048 × 2046 pixels, while 400 Hz reads out 5^◦^ × 1.25^◦^ at 2048 × 512 pixels. The lateral sampling, and hence *p*, is the same in both.

The pipeline has four stages (Figure 1). **(1) Preprocessing** (Section 3.2) removes edge artifacts, flags occluded (closed-eye) frames, and corrects the illumination profile. **(2) Registration** (Section 3.3) removes the global eye motion with a GPU patch-based translation stage and an optional elastix refinement. **(3) Postprocessing** (Section 3.4) forms a masked frame-difference stack that both renders the perfused vessel network and supplies the space– time images. **(4) Velocity measurement** (Section 3.5) traces a space–time image along each marked vessel and reads the velocity from the streak orientation with the Radon transform. The construction mirrors the AO retinal workflow of [1] so results are comparable, while contributing an open, GPU-accelerated registration front-end.

**Figure 1:**
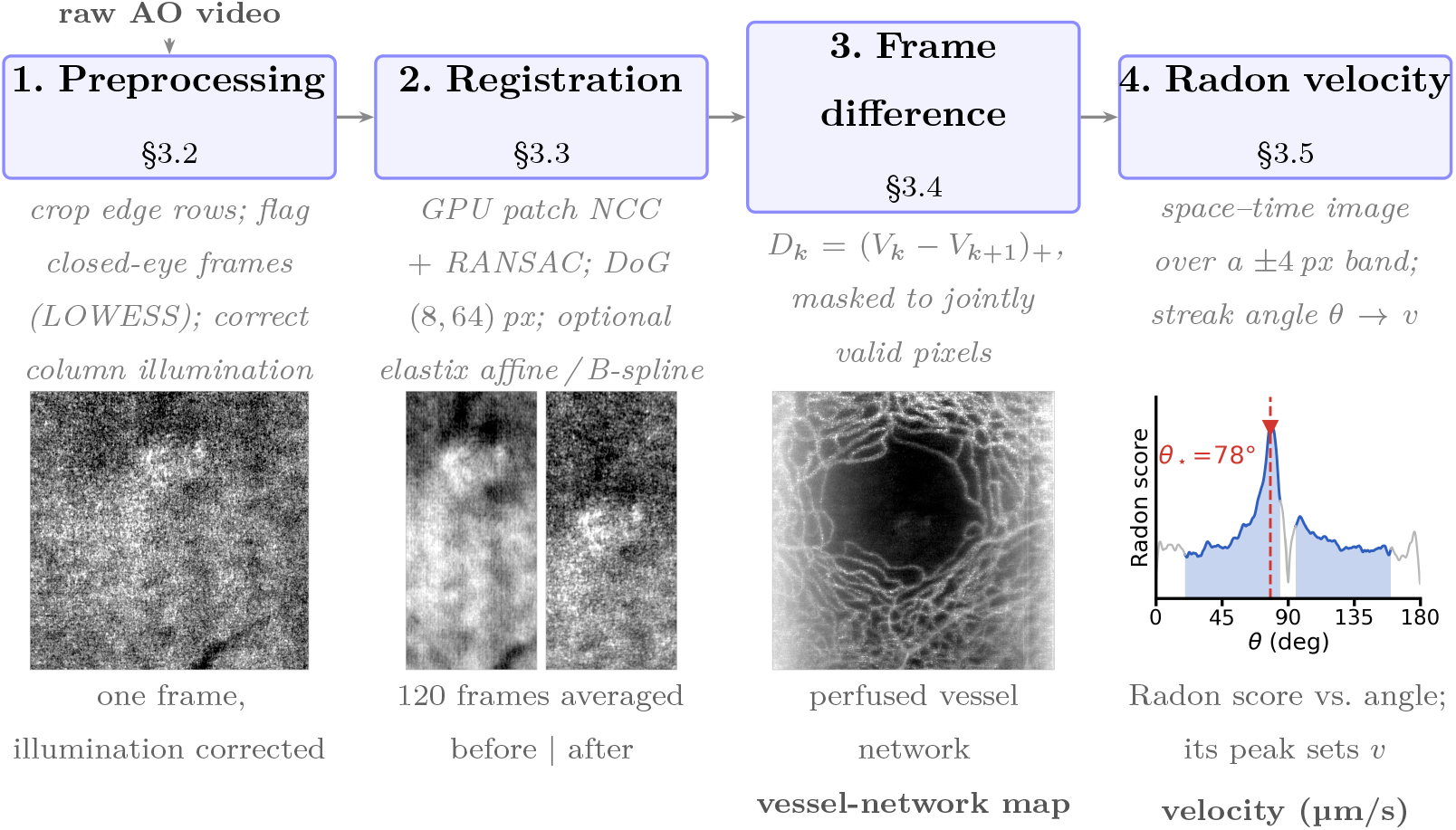
The four-stage pipeline, with the real data at each stage. Panels 1–3 are subject XC33, left eye, 100 Hz — the acquisition of Figure 5. **1.** Preprocessing crops the imager’s dark edge rows, flags closed-eye frames and removes the column illumination profile. **2.** Registration estimates a per-frame shift; the two tiles are the *same* 120 frames averaged the same way, with the stage disabled and enabled, so the difference between them is the stage’s effect alone — averaging without it smears the mosaic away (mean gradient magnitude rises 3.4× once motion is removed). **3.** Positive frame differencing cancels static tissue and leaves only moving blood; its masked mean projection is the perfused vessel-network map, with the foveal avascular zone dark at the center. **4.** A space–time image is traced along a marked vessel and the Radon transform scores every candidate streak angle; the peak of that score — here *θ_*_* = 78^◦^ within the [20^◦^, 85^◦^] search band (shaded) — sets the velocity through Equation (9). We plot the score rather than the space–time image because individual streaks are shallow and low-contrast, which is exactly why the angle is recovered statistically rather than by eye. This panel is the same window Figure 2 shows at full size (subject OT34, 400 Hz, the velocimetry mode).

The pipeline ships as two surfaces over one analysis core: an interactive 3D Slicer extension [33] for single videos, and a headless, JSON-driven batch tool for cohort-scale runs. Registration is implemented in PyTorch [34]; the elastix refinement uses the ITK Python bindings [6]; the velocimetry offers interchangeable CPU (scikit-image [35]) and GPU back-ends. Every stage is driven by a JSON configuration, and the preprocessing, registration, and postprocessing stages are idempotent (skipped when their output already exists), so an interrupted cohort run resumes cleanly. Volumes are exchanged as NRRD and tracings as Slicer markup files, so the two surfaces interoperate.

### 3.2. Preprocessing

The preprocessor runs four steps, in order. (1) *Edge-row removal*: the imager produces a few dark rows at the top of each frame; a fixed count is cropped per acquisition rate (80 rows at 30 Hz, 160 at 100 Hz, 90 at 200 Hz and 64 at 400 Hz) before any intensity statistic is computed. The rate is read from the file name, and an unrecognized rate is an error rather than a silent default. (2) *Occluded-frame detection*: we take each frame’s mean intensity, fit a smooth trend with a locally weighted smoother (LOWESS [36], span 0.3), and flag a frame as open only if its mean exceeds 0.75 of the trend; closed-eye frames are zeroed and excluded from later statistics. (3) *Column-wise illumination correction*: the illumination varies slowly across columns but is roughly uniform along rows, so we take the mean of the open frames as a one-dimensional column profile, smooth it with a boxcar 127 pixels wide at the native 2048-pixel width, normalize it to its peak, and divide each frame by it — with the divisor floored at 0.2, so no column is amplified by more than 5×. Columns at either edge whose profile stays below that same 0.2 are then trimmed. (4) *Smoothing and equalization*: an isotropic Gaussian blur (*σ* = 1 pixel) is applied by default to suppress sensor noise; an optional contrast-limited histogram equalization (CLAHE [37]) is off by default, since registration applies its own vessel enhancement. The output is an NRRD [38] stack at the original bit depth, with closed frames zeroed and edge columns trimmed.

### 3.3. Registration

Removing the global eye motion leaves the flow of blood cells as the only systematic temporal signal. Registration runs in two stages: a fast GPU patch-based translation on every frame, followed by an optional elastix refinement (translation → affine → B-spline).

#### GPU patch-based translation

The first stage estimates a per-frame shift **d***_k_* ∈ ℝ^2^ relative to a reference frame. It is kept rigid (a single translation) because it must run on every frame at the full rate — up to ∼8000 frames for a 20 s acquisition at 400 Hz — and because most ocular motion within the working second is well approximated by a global shift. To boost the match, frames are optionally downscaled by 0.5 and band-pass filtered with a Difference-of-Gaussians (DoG) kernel [39] that keeps capillary-scale structure while removing slow illumination,

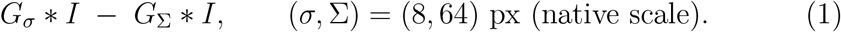

Reference patches (default size 449 × 449) are sampled on a grid; patches overlapping the field of view by less than 0.9 are dropped. For each patch we compute the normalized cross-correlation against the moving frame, evaluated efficiently as a GPU convolution with the patch kernels chunked across passes to fit memory. Each patch votes for the shift at its correlation peak, kept only if that peak is strong enough (correlation above 0.6, the active gate in our runs); an optional uniqueness test — the best peak well above the second-best — can further filter patches. The per-frame shift is then a RANSAC robust mean over the admitted patch votes [5], which ignores the outliers caused by specular highlights or blink edges; if RANSAC fails it falls back to a correlation-weighted mean. The reference frame is chosen automatically as the *centroid frame* — the wide-open frame whose shift is closest to all the others,

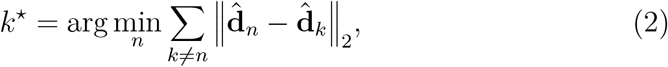

re-running the stage once if the initial guess was not the centroid. The candidate set here is narrower than the open frames of Section 3.2: *n* ranges only over frames whose mean intensity is above the LOWESS trend, not merely above 0.75 of it, so a partially occluded frame cannot become the reference; the sum over *k* still runs over every frame with a recovered shift. The initial guess is the first such frame. The stage emits the registered NRRD stack (shifts applied at integer-pixel offsets) and a per-frame displacement table.

#### Hierarchical elastix refinement

On the already-stabilized stack we optionally remove residual affine and non-rigid distortion with up to three elastix stages [6], run in sequence: translation, then affine, then a B-spline free-form deformation. Each stage is initialized by the *composition* of the previous transforms rather than by resampling the image between stages, which avoids cumulative interpolation blur.

The *affine* stage models a global linear warp about the image center **c**,

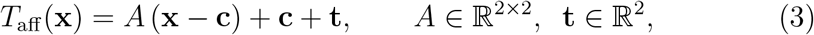

with six free parameters (the matrix *A* and translation **t**; the center **c** is fixed).

The *B-spline* stage adds a local free-form deformation [7]: displacements ***ϕ****_j_* on a regular control-point grid {**g***_j_*} of spacing *δ* (default 256 voxels) are blended by cubic B-spline kernels *β*^3^,

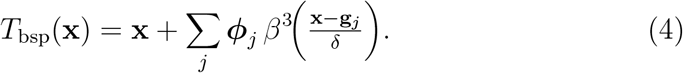

Both stages maximize the same similarity loss — the normalized mutual information [40, 41] between the fixed frame *I_F_* and the warped moving frame *I_M_*◦ *T*,

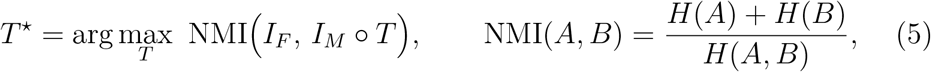

where *H*(·) and *H*(·, ·) are the marginal and joint image entropies; elastix minimizes −NMI by adaptive stochastic gradient descent [42] over 4096 random samples per iteration (up to 1000 iterations).

#### Multi-resolution

Each stage is solved coarse-to-fine over a four-level Gaussian image pyramid (down-sampling schedule 16, 8, 4, 1): the fixed and moving frames are blurred and down-sampled, registered at the coarsest level first, and the transform recovered there initializes the next finer level. This captures large displacements at coarse scales — where the similarity surface is smooth and free of spurious local optima — and refines fine detail at full resolution, which a single-resolution fit could not do without falling into a local minimum.

The B-spline stage is the most expensive, so it is gated by frame rate: by default it runs at 30 Hz and 100 Hz and is skipped above 150 Hz, that is at 200 Hz and 400 Hz (where a 20 s clip has 8000 frames). The gate is a switch, not a hard rule, and can be turned off. If a frame fails to converge (typically mid-blink while the reference is open), the stage falls back to the patch-based shift for that frame.

### 3.4. Frame-difference postprocessing

After registration, static structure (vessel walls, parenchyma, fixed noise) is stationary in pixel coordinates, while the only systematic motion left is blood flowing through the lumen. We exploit this by forming a *positive frame-difference* stack from the registered video *V_k_* (normalized to [*0, 1*]):

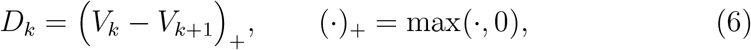

which keeps the moving-object signal and discards static content. Each difference is masked to the pixels valid in both frames (zero where registration left a blank border or an eye was closed), smoothed with an isotropic *σ* = 1 Gaussian, re-masked so that the blur cannot carry signal across the mask edge, and dropped if half or fewer of its pixels are valid. The blur is applied to the stack as a volume, so it acts along time as well as the two spatial axes and each difference is mixed slightly with its immediate neighbors. We save the difference stack together with three projections of it: a *masked mean difference*, which is the vessel-network map (Section 4.4); an unmasked mean difference, which differs only near the border, where the masked version compensates for pixels that were valid in only some frames; and a standard-deviation projection that also lights up every vessel carrying flow. A masked temporal mean of the registered video itself is saved alongside them as a high signal-to-noise-ratio (SNR) anatomical reference. Differencing helps for three reasons: it high-pass filters slow illumination drift; it cleanly separates flowing vessels (non-zero difference) from static ones (near-zero); and it produces the streak signal the Radon stage needs directly, with no separate detrending.

### 3.5. Velocity measurement using the Radon transform

We convert the frame-differenced stack into a per-vessel, per-time velocity in µm*/*s with a Radon-transform pipeline following the line-scan tradition [23, 24] and the flat-field angle estimator of Duncan et al. [8].

#### Space–time image

The user marks a vessel by placing control points along its center-line. The curve is resampled to one sample per pixel of arc length. For each frame *t* and curve sample *s* we take a maximum-intensity projection across a perpendicular band of ±*a* pixels (default *a* = 4) around the curve,

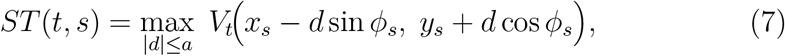

where *ϕ_s_* is the local tangent angle. Stacking all frames gives the *space–time matrix ST*. A cell moving along the vessel appears as a bright sloped streak: a fast cell moves many pixels per frame and so its streak is shallow; a slow cell’s streak is steep; a static feature is a vertical column.

#### Radon transform and two estimators

The Radon transform projects *ST* onto every line of orientation *θ*,

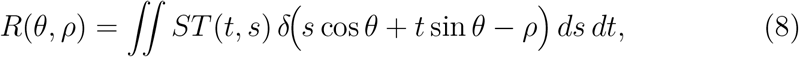

discretized over *θ* ∈ [0^◦^, 180^◦^) at *N_θ_* = 3600 angles (Δ*θ* = 0.05^◦^). The transform is taken over the whole rectangular window: scikit-image’s default restricts it to the inscribed circle, which would reduce a 64-frame × 32-px window to its central 32 × 32 block, and on real data the whole window is measurably more repeatable (Section 5). The streak orientation *θ_*_* shows up as a sharp ridge (Figure 2). The *max-projection* estimator reads the ridge as the per-angle maximum of *R*; it is fast but sensitive to isolated bright rows (e.g. a stray reflection) that smear across all angles. The *flat-field/SNR* estimator — our canonical one, implementing Duncan et al. [8] — first divides *ST* by the rank-one (separable) illumination model built from its row and column means, subtracts the residual mean, and then scores each angle by its root-mean-square energy normalized by the global sinogram energy, which is far more robust to those spikes. For either estimator the grid angle with the largest score is refined below the grid step by fitting a parabola through the score at that angle and at its two grid neighbours, with the shift clipped to half a step.

**Figure 2:**
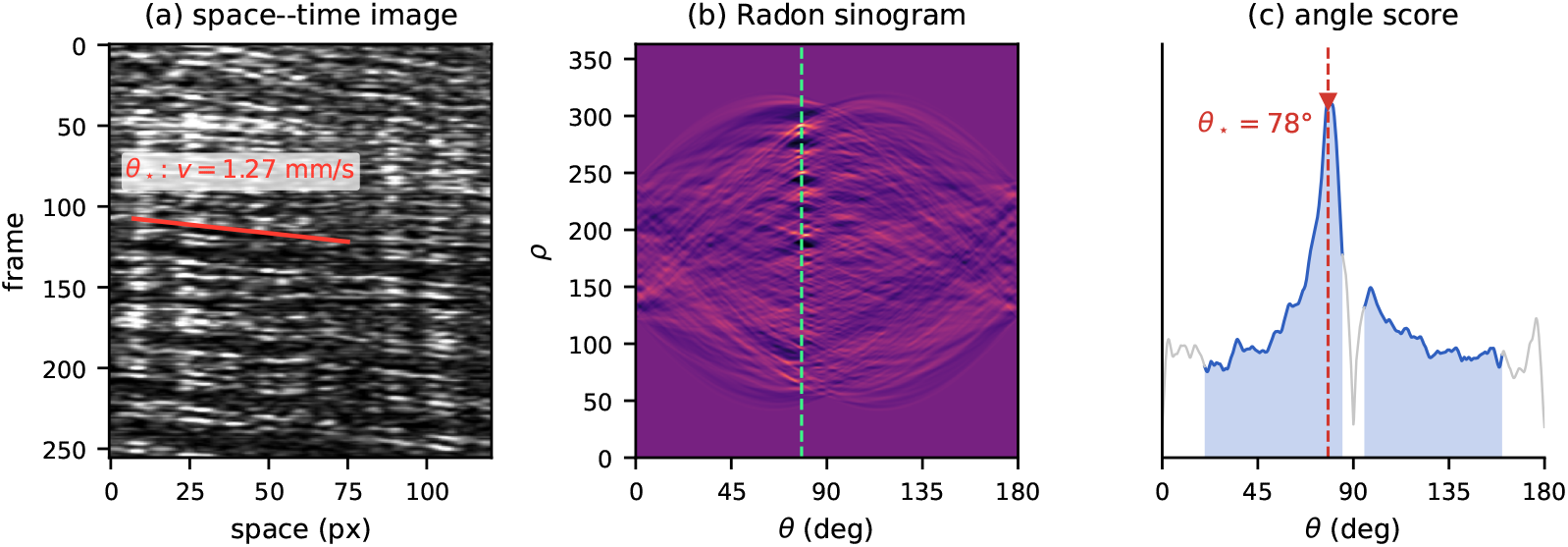
The velocity measurement, end to end, on one real capillary (subject OT34, 400 Hz, 256-frame window). **(a)** The space–time image: erythrocytes advance along the vessel between frames and so trace sloped streaks. The red line is the *measured* orientation, not a guide — it is drawn at the angle the estimator returned for this window, *v* = 1.27 mm/s. Note how shallow it is: at these velocities a cell crosses the traced segment in about 25 frames, so the streaks sit only ∼12^◦^ off the time axis and are individually low-contrast against the frame-to-frame intensity banding. **(b)** The Radon transform concentrates that orientation into a ridge at *θ_*_* (dashed). **(c)** The flat-field/SNR score over the search band [20^◦^, 85^◦^] (shaded; gray is excluded). Its peak is sharp — 2.6× the in-band mean — even though no single streak is obvious by eye, and that is precisely why the angle is recovered by integrating over the whole window rather than by tracing individual cells. The peak angle gives the velocity through Equation (9).

#### Angle-to-velocity

Both estimators share the same angle-to-velocity conversion; they may select slightly different optima *θ_*_* (Section 4.5). The streak slope ties the per-frame pixel displacement to the per-frame time 1*/f* (with *f* the true frame rate), so the physical velocity is

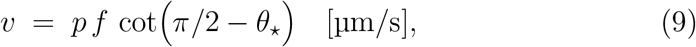

where *p* is the pixel pitch. The pitch enters *linearly*: an estimator that reports pixels per frame (effectively *p* = 1) is wrong by a factor 1*/p* — about 1.5× at the clinical *p* = 0.67 µm*/*pixel. We search *θ_*_* within a default band [20^◦^, 85^◦^] (excluding near-parallel and near-perpendicular streaks that map to ambiguous or infinite speeds), in the forward or reverse direction, and optionally constrain the search by physical velocity bounds (default 0 to 3000 µm*/*s, the physiological range of parafoveal capillary flow).

#### Sliding window

A single angle over the whole *ST* gives one velocity per vessel and loses the pulse. We instead run the estimator on windows of 64 frames, processed in parallel, dropping any window that contains a blank frame — one whose pixels sum to less than 5 intensity units along the vessel, which after preprocessing means a closed-eye frame. A window’s velocity is the dominant streak slope over all 64 of its frames, so a window that straddles a blink is reading the eyelid margin or the tear film sweeping across the field rather than blood; requiring every frame to be open is what makes the window a measurement of flow at all (Section 4.6). The resulting series is linearly interpolated to one value per frame and low-pass filtered with a fourth-order Butterworth filter at 10 Hz (default), run forward and backward so it is zero-phase, which removes window-boundary jitter while preserving heart-rate pulsatility (1–3 Hz). Long vessels can additionally be sliced into spatial windows (batch default 64 pixels; the Slicer module still defaults to 32) to study flow heterogeneity along the vessel. Both the batch tool and the Slicer module default to a temporal stride equal to the window length, so that windows do not overlap; every run reported here instead used a stride of 16 frames (a four-fold overlap), which is what run_all_velocity_cpu.sh passes on the command line.

## 4. Experiments

The quantitative core is the accuracy of the Radon velocimetry against synthetic ground truth (Section 4.5). We then check the registration’s inter-frame self-consistency on real video (Section 4.2), benchmark the translation stage’s speed and robustness against itk-elastix (Section 4.3), show the vessel-network map qualitatively (Section 4.4), report velocities recovered from real capillaries against the published values of Gu et al. (Section 4.6), and settle which Radon backend is canonical (Section 4.7).

### 4.1. Datasets and setup

#### Real video

The full registered cohort comprises 2344 acquisitions from 57 subjects, spanning 30 Hz to 400 Hz (parafoveal capillaries, *p* = 0.67 µm*/*pixel), and registration quality is evaluated across all of it (Section 4.2); the 100 Hz near-full-field subset renders the vessel network and the 400 Hz subset resolves erythrocyte motion for velocimetry. The velocity analyses here demonstrate the pipeline on three representative 400 Hz acquisitions from three subjects^1^ — OT34 (3336 frames), XC02 (3993 frames), and XC01 (3658 frames, counting in each case the post-difference frames that enter the velocimetry) — traced with 10 vessel curves in total (3, 3, 4), each split into 2–6 spatial sub-columns, giving 40 curve–column series. Sub-columns of one vessel are correlated, so for these three acquisitions the replication is 3 acquisitions (3 subjects) and 10 vessels; we keep this nesting explicit below.**[TODO: Per-subject demographics (the registration cohort spans 2344 videos from 57 healthy subjects).]**

#### Vessel-network video

Rendering the network uses the 100 Hz near-full-field acquisitions of the cohort; Section 4.4 shows one such map (subject XC33, lef eye).

#### Synthetic video

For velocity validation we use synthetic space–time videos with a known per-frame velocity, each 3 s at 400 Hz: *constant* (1200 µm*/*s), *linear* (ramp 500 µm*/*s to 4097 µm*/*s, mean 2298.5), and *pulsatile* (60-bpm sinusoid in 500 µm*/*s to 2000 µm*/*s, mean 1250).

#### Hardware and settings

Registration runs on the GPU (PyTorch); the velocimetry CPU backend uses scikit-image and the GPU backend PyTorch. Experiments ran on a workstation with an AMD Ryzen 9 5900X CPU (12 cores/24 threads) and 128 GB RAM. All timing results (Table 3, Figure 4c) were measured with an NVIDIA RTX A6000 GPU (48 GB), as was the synthetic capture-range sweep (Figure 4a). The real-frame head-to-head that supplies the similarity and failure-rate columns of Table 3 was measured on an NVIDIA RTX A1000. Only the timings are hardware-sensitive: re-running the GPU method on the A6000 reproduces all 800 stored per-frame shifts (largest difference 0.0000 px against a 0.01-px tolerance), so every similarity and failure figure is device-independent, and all reported timings are the A6000 figures. Unless stated otherwise the velocity backend is the CPU flat-field/SNR estimator (the GPU backend returns the same velocities, Section 4.7) with *θ* ∈ [20^◦^, 85^◦^], *N_θ_* = 3600, a 64-frame temporal window, 32-pixel spatial windows (a deliberate override of the batch tool’s 64-pixel default, to resolve intra-vessel variation), and a 10 Hz Butterworth post-filter.

**Table 2:**
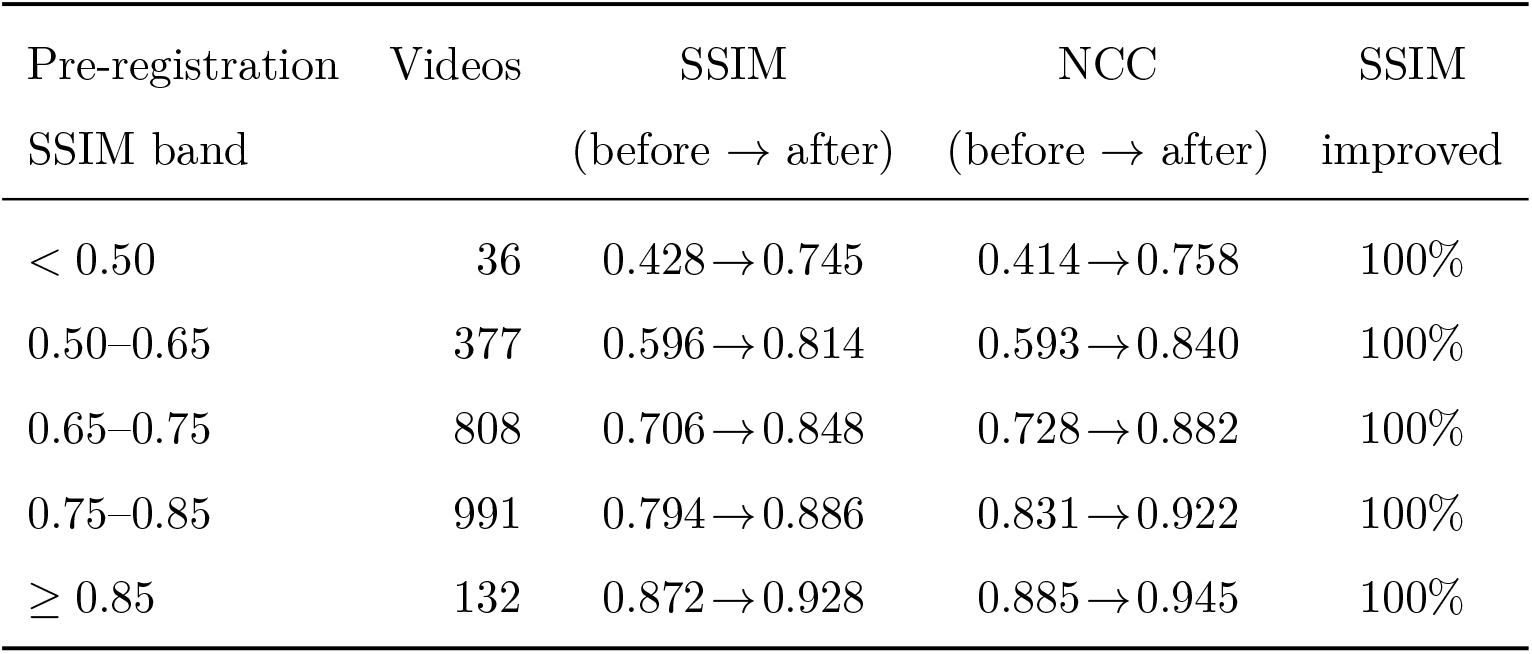
Registration gain stratified by input quality, over the same 2344 acquisitions as. **Table 1**. Acquisitions are banded by their pre-registration (re-shift proxy) SSIM, so the top row holds the worst-quality acquisitions in the cohort. Registration improves SSIM on every acquisition in every band, and the mean gain is largest where the input is worst.

**Table 3:** Per-frame registration time and post-alignment quality over the benchmark set — 20 acquisitions drawn from the registered cohort of. **Table 1 and re-registered from scratch for this comparison (10 at 100 Hz from 10 subjects, and 10 at 400 Hz all from a single subject imaged at two visits; 40 sampled open frames each) — patch-based translation versus itk-elastix.** †Time is the per-frame range from 400 Hz (small frame) to 100 Hz (large frame): the patch method is batched on one GPU, while elastix runs per-frame on 8 CPU threads (it offers no batched or GPU path; the thread count is our choice, not a limit of the library). SSIM/NCC are the median image similarity of each *translation-aligned* frame to its reference (no ground-truth shift exists on real data); a failure is a missing shift, one exceeding 1.2× the frame’s largest dimension, or an aligned SSIM below 0.30 — on these data only the last rule ever fires, and the ranking is unchanged for any floor from 0.20 to 0.45. Timings are from the RTX A6000; the similarity and failure columns were measured on an RTX A1000; re-running the shift estimation on the A6000 reproduces all 800 of them exactly. The same tuned pyramid with a mutual-information metric runs 2.2× slower than the normalized-correlation one and was not scored on the real frames, so it is omitted here; its synthetic behavior is in the text — better under occlusion, worse in capture range and under illumination gradients.

| Method | Time/frame<sup>†</sup> | SSIM ↑ | NCC ↑ | Fail. % ↓ |
| --- | --- | --- | --- | --- |
| Patch-NCC + RANSAC (ours, GPU) | 0.06 s to 0.73 s | **0.385** | **0.638** | **9.8** |
| elastix transl., default 4-level (CPU) | 3.6 s to 4.0 s | 0.317 | 0.402 | 38.8 |
| elastix transl., tuned 6-level NCC (CPU) | 2.3 s to 2.7 s | 0.353 | 0.562 | 25.9 |

#### Software environment

Linux, Python 3.13.9, NumPy 2.2.6, SciPy 1.16.3, pandas 3.0.1, PyTorch 2.6.0 (CUDA 12.4), scikit-image 0.25.2, scikit-learn 1.8.0, statsmodels 0.14.6, and itk-elastix 0.25.3 (ITK 5.4.6). These are the versions of the workstation that produced every result below except Table 1, not merely the latest the code accepts; a frozen listing of its whole environment ships with the code. The cohort registrations and the similarity metrics of Table 1 were computed by a co-author on a separate workstation with the released pipeline and evaluator; the package versions there were not recorded.

### 4.2. Registration quality

Because the unregistered stacks were not retained for most of the cohort, we report an internal *self-consistency* check here rather than a before/after against the raw frames; the head-to-head against a competing method, on the 20 acquisitions whose unregistered frames were kept, is Section 4.3. For each acquisition we sample up to 120 evenly spaced pairs of consecutive eye-open frames (six short videos supply fewer, at least 8) on a central 1024 × 1024 crop, clipped to the 934 and 448 rows that remain after edge trimming at 200 and 400 Hz, and compute four image-similarity scores — structural similarity (SSIM [43], higher better), normalized mutual information (NMI, higher better), normalized cross-correlation (NCC, higher better), and normalized RMS error (NRMSE, lower better) — comparing the registered stack (“aligned”) against the same pairs with the recorded inter-frame shift re-injected (“re-shifted”, the “before”). The re-shifted proxy carries a one-sided interpolation/border penalty, so the gap overstates the pure registration effect; with that caveat, Table 1 shows registration improves alignment consistently across the whole registered cohort (2344 videos from 57 subjects, spanning 30 Hz to 400 Hz) — raising SSIM from 0.73 to 0.86 and NCC from 0.75 to 0.89 on average, at every acquisition rate. SSIM and NMI improve on every one of the 2344 videos, and 2334 improve on all four metrics (the remaining 10 show an NCC decrease or NRMSE increase, all under 0.09; the largest is an NRMSE increase of 0.080, on a 400 Hz acquisition). The 30 Hz row rests on only 10 acquisitions from 5 subjects and is reported for completeness rather than as a characterization of that rate.

The corrected motion is large across the cohort (Figure 3): the per-video median per-frame shift averages 132 pixels (per-rate averages 89–137 px) — i.e. tissue motions of roughly 60 µm to 92 µm per frame — confirming that whole-frame stabilization is essential before any per-pixel analysis. On average 93% of frames per acquisition are eye-open and usable (median 96%).

**Figure 3:**
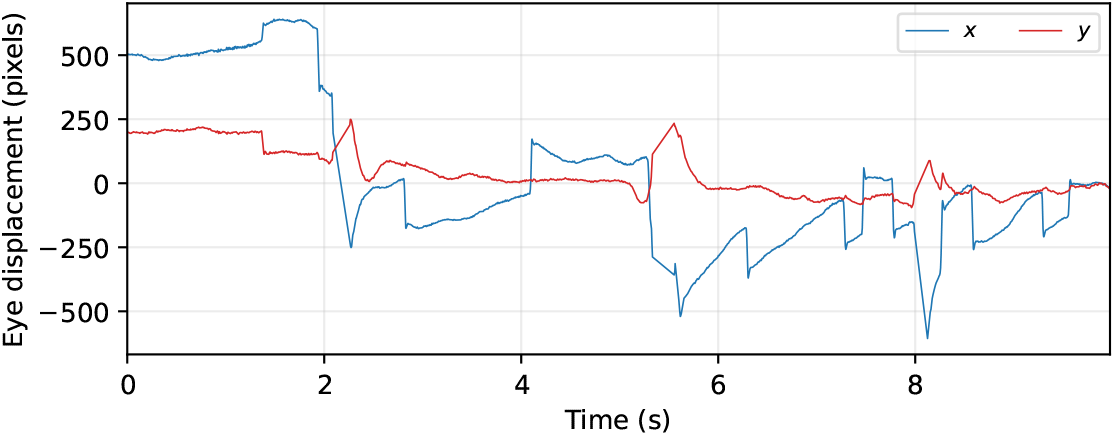
Recovered per-frame eye displacement from the patch-based stage for a representative acquisition (subject XC02, 400 Hz). The trajectory spans hundreds of pixels and is dominated by slow drift punctuated by micro-saccades.

#### Video quality and screening

Acquisition quality varies widely in a cohort of this size, so we state the screening explicitly. Two automatic rules act, both defined in Section 3.2: closed-eye frames are detected per frame and excluded from every statistic, and an acquisition is dropped entirely if the pipeline cannot produce a registered stack from it (3 of 2347 staged acquisitions, leaving the 2344 reported here). Beyond that, *no acquisition was excluded on subjective image quality*. That is deliberate: the claim being tested is that registration works across the full range of video the instrument produces, and pre-selecting good video would make the claim circular. The retained range is genuinely wide — pre-registration SSIM spans 0.14 to 0.97, and the eye-open fraction spans 0.6% to 100% (median 96%).

Two checks show the headline result is not carried by the good acquisitions (Table 2). First, stratifying by pre-registration similarity, registration improves SSIM on 100% of acquisitions in *every* quality band, and the improvement is largest exactly where the input is worst (+0.32 in the lowest band against +0.06 in the highest) — the expected behavior, since a well-aligned acquisition has less to gain. Second, imposing a hard quality floor changes nothing material. Requiring an eye-open fraction of at least 0.50 removes 16 acquisitions (0.7%) and moves no cohort mean in Table 1 by more than 0.0007; a floor of 0.80 removes 114 (4.9%) and moves none by more than 0.003, the largest being post-registration NCC (0.894 → 0.896). The before-to-after *gain* is steadier still: under either floor no gain in Table 1 shifts by more than 0.0014. Screening therefore makes registration look marginally better, not worse, so we report the unscreened cohort as the conservative choice and note that a deployment wanting a quality gate can apply one at no cost to these numbers.

### 4.3. Registration speed and robustness versus itk-elastix

The patch-based translation stage runs on every frame at the full acquisition rate, so it must be both fast and robust to the large eye motions of high-speed AO video. We benchmark it directly against the standard gradient-based alternative — the translation transform of itk-elastix [6], optimized by adaptive stochastic gradient descent over a multi-resolution image pyramid — estimating a per-frame shift on the same frames, with neither method seeding the other. We compare the baseline in three configurations: elastix with its default four-level pyramid, and a hand-tuned coarse six-level pyramid with either a mutual-information or a normalized-correlation metric.

All three use our pipeline’s own metric and optimizer budget rather than the library’s stock values — normalized mutual information, 1000 iterations and 4096 spatial samples, against stock defaults of Mattes mutual information [44], 256 iterations and 2048 samples — so the baseline is given a 4× larger optimizer budget and 2× the sampling of an untouched elastix install. The optimizer itself (adaptive stochastic gradient descent) is left at its default.

#### Capture range (synthetic ground truth)

We crop shifted windows of real frame content (no zero-fill borders, which would artificially penalize an intensity metric) with a *known* integer shift and recover it; Figure 4(a) plots the fraction recovered within 5 px over 32 trials per displacement (four reference frames × eight random directions). The patch method recovers *every* shift across the full 5–300 px range (median error 0.28 px). Default-pyramid elastix holds 100% only to ∼30 px, falling to 97% at 50 px, 66% at 120 px and 0% at 300 px; the tuned six-level pyramid holds at least 97% out to ∼170 px and then diverges sharply by metric — at 300 px the normalized-correlation configuration still recovers 69% while the mutual-information one recovers 12%. The patch method needs no capture-range tuning because its correlation search is global. Neither method degrades appreciably with additive noise (every configuration scores ≥ 97% at every level). Under slow illumination gradients the patch method holds 100% throughout, whereas the default and mutual-information elastix configurations fall to 59% and 69% at the strongest gradient (tuned normalized-correlation holds 91%). That axis applies one smooth full-field linear ramp to the moving frame, so it exercises a frame-to-frame illumination difference, not a localized band travelling across a sequence — which is the perturbation the velocimetry is actually sensitive to (Section 4.6). The occlusion, noise and illumination sweeps carry a fixed 40 px baseline offset, and every trial on every axis carries light baseline noise (*σ* = 0.02, replaced by the swept value on the noise axis), so a configuration’s residual failures reflect those as well as the perturbation being swept. Every percentage on these four axes is a 32-trial estimate, and cells in the middle of a method’s transition from success to failure carry the most trial-to-trial variability: repeating the whole sweep under an independent seeding moved 87 of the 88 cells by at most 10 points — the exception being the patch method at a 30% occluding block, its breakdown point, which moved by 28 — and changed the rank order of the four configurations at no perturbation level beyond ties at 100% broken by a single trial. Both sweeps are released, and make_figures.py prints this comparison.

**Figure 4:**
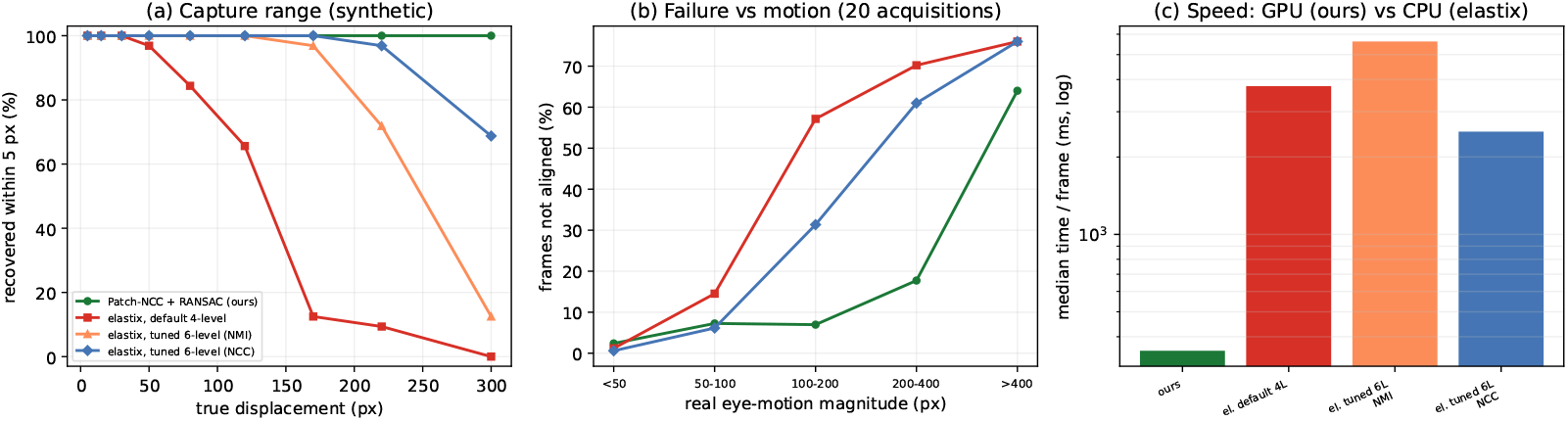
Registration benchmark, patch-based translation versus itk-elastix. **(a)** Capture range on synthetic ground truth: fraction of known shifts recovered within 5 px (32 trials per displacement). The global patch search holds 100% to 300 px while default-pyramid elastix degrades past ∼30 px and even the tuned pyramid past ∼170 px. **(b)** Benchmark set (20 acquisitions): per-frame failure rate versus eye-motion magnitude — the patch method’s advantage grows with motion. **(c)** Per-frame compute time of the registration core (log scale): the GPU patch method versus CPU elastix.

#### Speed

Timed with the patch method batched on one RTX A6000 GPU and elastix per-frame on 8 CPU threads (elastix offers no batched or GPU path; the thread count is our choice, not a limit of the library), the patch method registers a 100 Hz frame in 0.73 s and a 400 Hz frame in 60 ms, against 2.3 s to 6.0 s per frame for elastix — a speed-up of ∼3.6× (100 Hz) to ∼38× (400 Hz) over the *fastest* elastix configuration, and larger over the others (Figure 4c, Table 3). The gap widens at 400 Hz because the smaller frames let the patch method batch many frames per GPU pass while elastix stays sequential.

Three qualifications make these an upper bound on the practical advantage. First, they are registration-*core* times: the patch method’s band-pass and half-resolution rescale — a CPU step costing 219 ms per 100 Hz frame and 53 ms per 400 Hz frame on this workstation — sit outside the timed core, whereas the elastix times include everything elastix does. Charged end-to-end the speed-up is ∼2.8× (100 Hz) and ∼20× (400 Hz). Second, the patch stage searches at half resolution and so processes 4× fewer pixels than elastix, which is part of why it is faster. Third, we did not attempt a throughput-matched CPU baseline running one single-threaded elastix process per core, which would narrow the gap further. The timing runs cover two acquisitions per rate.

#### Benchmark on real frames

On the benchmark set (10 100 Hz + 10 400 Hz acquisitions, 40 sampled open frames each) we register every frame to its reference from scratch and score the post-alignment image similarity. Each method’s recovered shift is applied the same way — at its full sub-pixel value, by bilinear resampling, over the region the two frames still share — so the comparison comes from the shift estimates alone and not from a resampling policy. (The production pipeline instead truncates the shift to whole pixels and resamples with nearest-neighbor, which is why the registered stacks it writes carry no interpolation blur; that choice is common to all methods and would not change the ranking.) A frame counts as a failure if a method returns no shift, one exceeding 1.2× the frame’s largest dimension, or leaves the aligned SSIM below 0.30 (Table 3). On these data only the last rule ever fires — no method returned a missing or implausibly large shift — so the reported rates are governed entirely by that floor. The ranking is insensitive to where the floor is put: the patch method has the lowest failure rate at every threshold we tested from 0.20 to 0.45 (0.2% vs 2.5–5.9% at 0.20; 9.8% vs 25.9–38.8% at 0.30; 35.1% vs 47.9–64.2% at 0.35), though the absolute rates naturally are not, which is why we also report the threshold-free median SSIM and NCC. The patch method leaves the fewest frames unaligned (9.8%, vs 38.8% default and 25.9% tuned-NCC) and gives the highest aligned similarity. The advantage grows with eye motion (Figure 4b): for frames moving 100 px to 200 px — the benchmark-set median is 115 px — the patch method fails on 7% versus 31–57% for elastix; only beyond 400 px (rare, mid-saccade) do all methods struggle (the patch method at 64%).

#### Where elastix is competitive

Two caveats keep the comparison honest. First, within its capture range elastix is slightly more accurate (sub-pixel, vs the patch method’s half-resolution ∼ 0.5 px). Second, on the 1024×1024 synthetic field — a quarter of a full 100 Hz frame — with a large (≥ 30%) opaque occluding block, the default and mutual-information elastix configurations stay at or above 97% recovery whereas the patch method can be starved of un-occluded patches (59% at a 30% block, 6% at 40%); this did not surface on real frames (above), whose larger field yields many more patches, but it bounds the patch method when the usable image area is small. The comparison is not uniform across elastix configurations, though: the tuned normalized-correlation pyramid — the strongest elastix baseline everywhere else — is the most occlusion-sensitive of all four methods, at 0% and 3% on the same blocks.

### 4.4. Vessel-network map

Projecting the masked frame-difference stack (Section 3.4) over time renders the perfused capillary network: vessels carrying moving cells have a fluctuating, non-zero difference signal and appear bright, while static tissue cancels — the same motion-contrast principle used by [1] and, in other modalities, OCTA [2]. Figure 5 shows the network from a 100 Hz near-full-field acquisition (subject XC33), the rate dedicated to wide-field network mapping; the parafoveal capillary arcades and the foveal avascular zone are clearly resolved.

**Figure 5:**
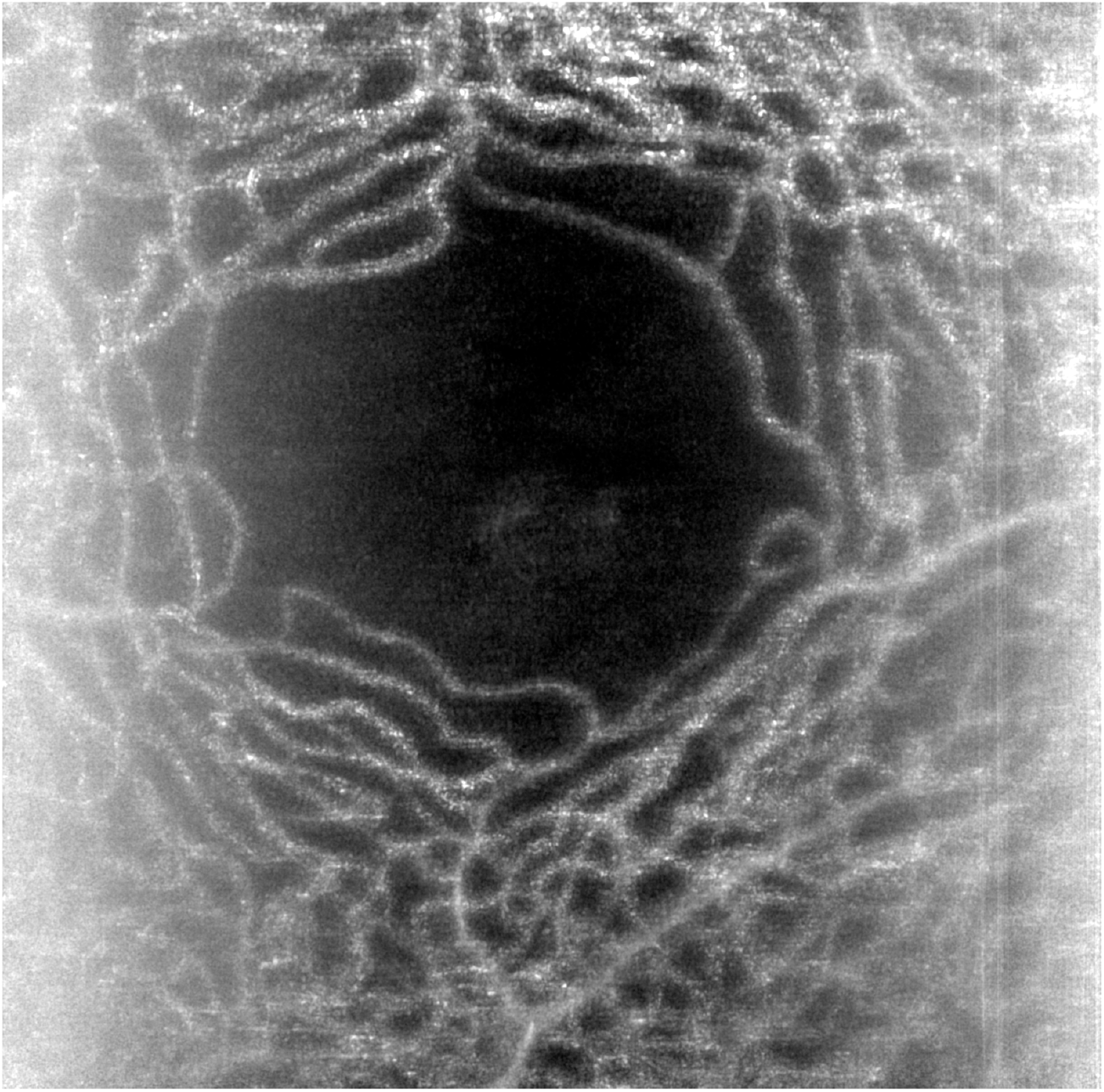
Perfused parafoveal capillary network rendered as the temporal mean of the masked frame-difference stack from a 100 Hz near-full-field acquisition (subject XC33, left eye). Vessels carrying moving erythrocytes appear bright while static tissue cancels; the dark central region is the foveal avascular zone. This is the pipeline’s wide-field network-mapping mode (cf. the 400 Hz velocimetry mode used elsewhere).

### 4.5. Radon velocimetry accuracy on simulation

Table 4 reports the velocity recovered from the three synthetic regimes for both estimators, in single-window and sliding-window (“series”) modes; the ratios are the recovered series *mean* over ground truth. Single-window mode applies one Radon transform to the whole 1200-frame space–time image; series mode is the tool’s default, 64-frame windows at a 16-frame stride. In series mode the flat-field/SNR estimator recovers the mean to within ±4% on all three regimes, and max-projection to within ±5%. One window over the whole acquisition is a different matter: on constant flow it agrees with the series (0.997× flat-field/SNR, 0.958× max-projection), but on the non-stationary regimes it fails, because the streak angle changes with time and one sinogram superimposes every slope in the video, so neither metric has a single ridge to find — flat-field/SNR over-estimates (1.301× on linear, 1.485× on pulsatile) and max-projection returns a small fraction of the true velocity (0.063×, 0.126×). That is why the tool measures in sliding windows, and every other velocity in this paper is a series-mode result. Between the two estimators, flat-field/SNR is the more stable: across the seeded replicates reported below its across-seed spread is the tighter on every regime. Mean accuracy is not instantaneous accuracy: the per-window scatter is tiny on constant flow (SD ≈ 3 µm*/*s to 27 µm*/*s) but large on the non-stationary regimes (41–46% of the mean for either estimator), which is signal, not noise, and bounds how finely one window can resolve the waveform. This benchmark isolates the estimator (a straight vessel, no eye motion, direct center-line sampling), so it validates the angle-to-velocity calibration but does not exercise registration or the curved-vessel space–time construction.

**Table 4:**
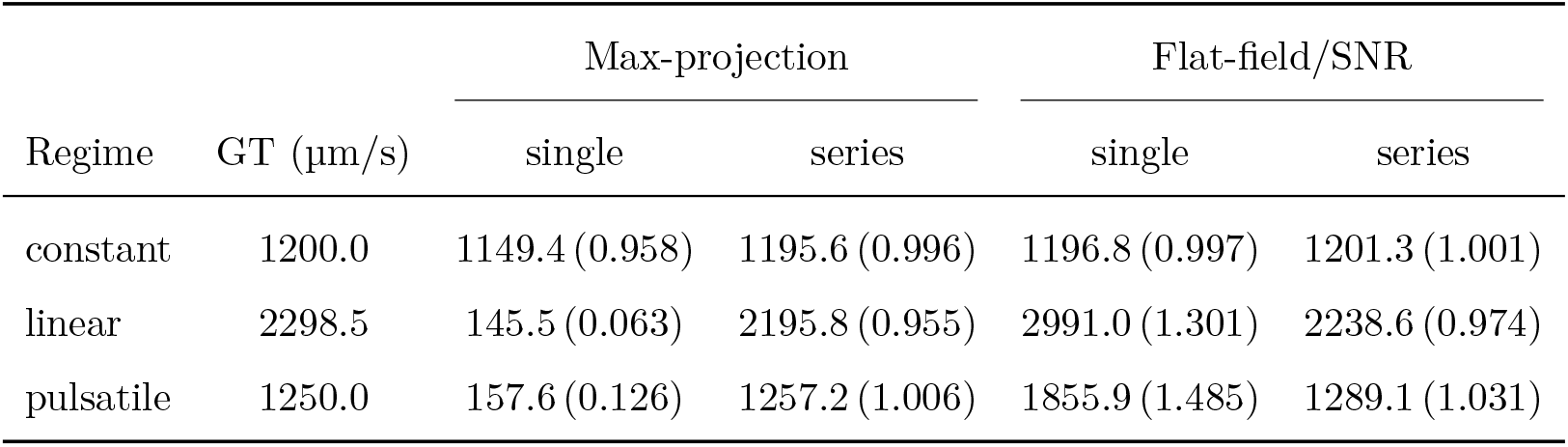
Recovered vs. ground-truth *mean* velocity on synthetic flow (ratio estimate/GT in parentheses; one video per regime, with 20-seed replicates reported in the text). Single mode is one Radon transform of the whole 1200-frame space–time image; series mode uses a 64-frame sliding window and is the mode the tool ships. The per-window SD in series mode (flat-field/SNR / max-projection) is 3.0*/*26.6 (constant), 1013*/*996 (linear), and 535*/*525 µm*/*s (pulsatile): instantaneous scatter is much larger than the mean-accuracy ratio on non-stationary flow.

One video per regime leaves the ratios open to the objection that they are a lucky draw, so we repeated the whole benchmark on 20 independently seeded videos per regime (run_replicates.py; Table 4 is seed 0). Across the 20 seeds the series-mode ratio is 1.001_±_._007_ (max-projection) and 1.000_±_._001_ (flat-field/SNR) on constant flow, 0.963_±_._023_ and 0.968_±_._003_ on linear, and 0.991_±_._018_ and 1.028_±_._014_ on pulsatile, where the subscript is the across-seed sample standard deviation, as in Table 1. Every regime-and-estimator mean sits within 4% of ground truth, so the accuracy figure characterizes the estimator and not the particular video. Individual videos are less forgiving: the worst single seed is off by 11.1% (max-projection, linear) and by 5.1% for the flat-field/SNR estimator (pulsatile), which is why the accuracy claim is made for series-mode means rather than for one window or one acquisition. The published ratios are not systematically flattering — the seed-0 linear flat-field/SNR ratio (0.974) is nearer unity than the 20-seed mean (0.968), but the seed-0 constant max-projection (0.996) and pulsatile flat-field/SNR (1.031) ratios are farther from unity than their own means (1.001 and 1.028). Because *v* = *p f* cot(*π/*2 − *θ_*_*), the pixel pitch enters linearly. We verified this on the constant-flow synthetic at the clinical pitch *p* = 0.67 µm*/*pixel: the flat-field/SNR estimator — the one used everywhere else in this paper — recovers 1205 µm*/*s in series mode, within ±2% of the 1200 µm*/*s ground truth, and over 20 seeds 1.004_±_._002_ with a worst single seed of 0.7%. Max-projection is looser at this pitch (1.013_±_._017_, worst seed 5.6%), consistent with its sensitivity to transient bright rows. A velocimeter that omits *p* woul over-estimate by 1*/p* ≈ 1.49, more than an order of magnitude larger than either estimator’s error. All velocities in this paper use the calibrated formula (Figure 6).

**Figure 6:**
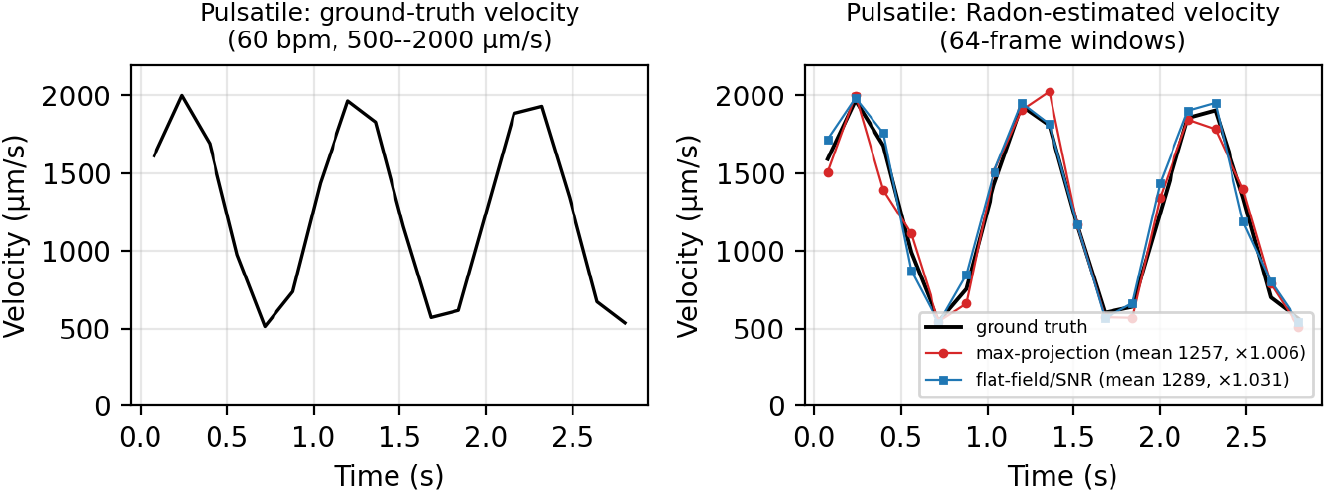
Radon velocimetry on the synthetic pulsatile regime: the ground-truth velocity waveform (left) and the Radon-recovered velocity from both estimators (max-projection and flat-field/SNR) against ground truth (right). The estimators track the 60-bpm waveform in amplitude and phase.

### 4.6. Erythrocyte velocity in real capillaries

This subsection is an end-to-end feasibility demonstration on real data: with three subjects it shows the pipeline produces physiologically plausible, pulsatile velocities, but it is not a population-level velocity validation (for which a far larger cohort and an independent in-vivo reference would be needed). We applied the canonical CPU flat-field/SNR velocimetry to the 40 curve–column series. For each we report the mean velocity *V_m_*, a robust systolic estimate *V*_max_ (95th percentile), a robust diastolic estimate *V*_min_ (5th percentile), and the pulsatility index PI = (*V*_max_ − *V*_min_)*/V_m_*. Because sub-columns of one vessel are correlated, we report per acquisition (*n* = 13*/*15*/*12) and pooled, and read the pooled ±SD as within-series dispersion, not a between-subject confidence interval (Table 5).

**Table 5:** Erythrocyte velocity in real parafoveal capillaries, per acquisition and pooled over 40 series (mean ± population SD, NumPy ddof= 0; the ± is within-series dispersion, not a between-subject CI), vs. the published AO values of Gu et al. [1]. For all 40 series the 95th percentile stays well below the *v*_max_ = 3000 µm*/*s search ceiling (the largest is 2.21 mm*/*s), so every *V*_max_ and PI here is a measured peak, not a right-censored one. Only windows whose 64 frames are all eye-open are measured; windows straddling a blink are dropped rather than smoothed over, which is what keeps the ceiling clear (Section 5).

| | $n$ | $V_m$<br>(mm/s) | $V_{\max}$<br>(mm/s) | $V_{\min}$<br>(mm/s) | PI |
| --- | --- | --- | --- | --- | --- |
| This work, OT34 | 13 | $1.03 \pm 0.11$ | $1.61 \pm 0.22$ | $0.50 \pm 0.18$ | $1.08 \pm 0.29$ |
| This work, XC02 | 15 | $0.92 \pm 0.11$ | $1.57 \pm 0.20$ | $0.37 \pm 0.10$ | $1.31 \pm 0.20$ |
| This work, XC01 | 12 | $0.90 \pm 0.15$ | $1.56 \pm 0.30$ | $0.27 \pm 0.13$ | $1.44 \pm 0.36$ |
| This work, pooled | 40 | $0.95 \pm 0.14$ | $1.58 \pm 0.24$ | $0.38 \pm 0.17$ | $1.28 \pm 0.32$ |
| [1], all vessels |  | 1.49 | 2.24 | 0.94 | 0.88 |
| [1], single-file |  | 1.22 | 1.88 | 0.76 | 0.91 |

The velocity search stops at *v*_max_ = 3000 µm*/*s, and no window reaches it. Of the 6328 eye-open windows behind Table 5 none returns 2974 µm*/*s or more — the band’s last grid angle, which the sub-grid refinement could carry half a step further to 2988 µm*/*s — and the largest single window velocity anywhere in the cohort is 2728 µm*/*s. Every series’ 95th percentile therefore sits well below the ceiling (the largest is 2.21 mm*/*s), so all 40 *V*_max_ and PI values are measured rather than right-censored.

That is a property of the window guard, not of the estimator, and it is worth being explicit about because it moved the table. Measured with the previous rule, which kept a window until 70% of its frames were blank, 204 of 7143 eye-open windows (2.9%) sat at the ceiling and one XC02 column was censored there. Every one of those windows straddled a blink, and not one of the windows whose 64 frames were all eye-open reached the ceiling at all: the estimator was reading the eyelid margin or the tear film sweeping across the field, which is broad and fast, instead of blood (Section 5). Requiring every frame of a window to be open drops 855 of 7586 windows (11%) and removes that population entirely. It also lowers the pooled *V*_max_ from 1.91 to 1.58 mm*/*s and the pooled PI from 1.55 to 1.28; XC02, the acquisition with the most blinks, moves most (*V*_max_ 2.26 → 1.57 mm*/*s, PI 1.84 → 1.31), and the pooled *V_m_* falls from 1.01 to 0.95 mm*/*s. The recovered mean velocities (*V_m_* ≈ 0.95 mm/s) are of the same order as, though somewhat below, previously reported AO parafoveal means (Gu’s single-file 1.22 and all-vessel 1.49 mm/s); they sit inside both of the diastolic-to-systolic excursions Gu reports (all-vessel 0.94 mm*/*s to 2.24 mm*/*s; single-file 0.76 mm*/*s to 1.88 mm*/*s). We do not claim a like-for-like match: vessel caliber differs between studies, and our diastolic *V*_min_ is lower and PI higher because our percentile estimate is not cardiac-gated and captures transient near-stall events (leukocyte passage, confluence interactions). We compare against Gu et al. alone because it is the matched instrument family and acquisition mode. The other AO capillary-flow measurements [29, 30] are of the same order — Tam et al. report plasma-gap speeds of 1.30 ± 0.55 mm/s across all capillaries and 1.80 ± 0.22 mm/s for leukocytes in leukocyte-preferred paths — but they report the speed of *individual* identifiable objects, in Tam et al. a leukocyte or a plasma gap traced across a space–time plot, in vessels selected for single-file flow (3.5 µm to 6 µm), rather than the dominant slope over a window of whatever the vessel carries. Tabulating those beside ours would invite exactly the like-for-like reading we are disclaiming. The recovered series are distinctly pulsatile (Figure 7).

**Figure 7:**
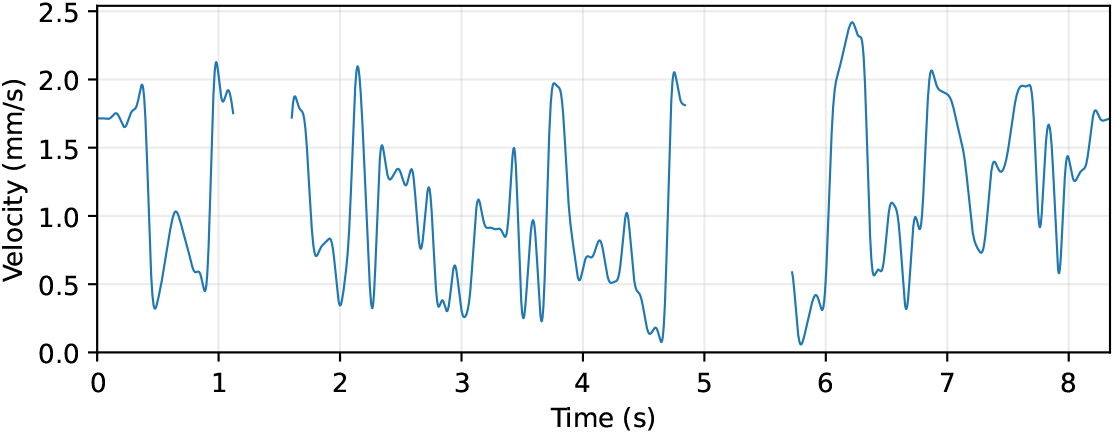
Recovered erythrocyte velocity waveform from one real parafoveal capillary (subject OT34, mean ∼1.12 mm/s), CPU flat-field/SNR estimator with a 10 Hz Butterworth filter. This series was chosen to show the waveform structure clearly, not at random: among the series with at least 55% eye-open coverage and under 1% of frames at the velocity-search ceiling (a condition every series now meets), it is the most pulsatile. Per-series statistics for the full set are in Table 5. The trace is distinctly cardiac-pulsatile; gaps correspond to blink/occlusion frames.

### 4.7. Backend consistency

The velocimetry offers CPU (scikit-image Radon) and GPU (PyTorch) backends. They share the flat-field normalization, the SNR metric, the band-limited peak search with its sub-grid refinement, and the angle-to-velocity conversion, and differ only in how the sinogram is computed: scikit-image rotates the zero-padded window and sums its columns, and the GPU backend performs the same rotation as a batched bilinear resampling of every window at every angle, padded to the same diagonal and rotated in the same sense. A standalone test checks the GPU sinogram against scikit-image’s on rectangular windows (to within 10^−4^ relative) and the two velocities on real streak windows. On the data behind Table 5 — every measured 64-frame window of the ten curves’ 32-px columns at the 16-frame stride, with the tool’s default search band, 6731 windows — the two backends return the same direction on every window and the same velocity to a median of 0.003 µm*/*s, with no window differing by as much as 1 µm*/*s (largest 0.8 µm*/*s) (compare_backends.py). The agreement is this tight only because the window guard is in force. Over the larger pre-guard window set the same comparison left eight windows (0.1%) differing by more than 1 µm*/*s, seven of them at the search ceiling: there the angle–velocity map is steepest, half a grid step of refinement is worth 15 µm*/*s, and the two sinograms’ 10^−4^ disagreement was enough to flip the refinement and the velocity by 14.6 µm*/*s (0.5%). The GPU backend processes the 6731 windows in 29 s against 3015 core-seconds on the CPU (0.45 s per window on one core, about 100×), so the CPU results reported here can be reproduced on the GPU without a change in the numbers beyond that tolerance. We keep the CPU backend canonical because its sinogram is scikit-image’s own and its calibration is the one Table 4 validated directly; the GPU backend inherits that validation through the agreement just described. An earlier GPU projector did not: it rotated the window in the opposite sense and disagreed with the CPU by up to 1.7× on real columns (Section 5).

## 5. Discussion

### Why the SNR estimator wins on pulsatile flow

On steady flow the two estimators pick almost the same angle in series mode: on the video in Table 4 they differ by 0.5%, and across the 20 seeded replicates their mean ratios differ by 0.1% — under 1% either way. Over one whole-video window they separate even on steady flow (3.9 and 8.6 percentage points), and it is max-projection that moves (0.958× against 0.997×): an isolated bright row (a transient reflection or a brief bright cell) projects onto *every* angle and gives the max-projection estimator a spurious flat ridge that biases its angle, whereas the RMS-per-angle metric averages those spikes away. On pulsatile flow neither estimator survives a whole-video window (1.485× and 0.126× in Table 4), because the streak angle changes with the cardiac phase and one sinogram superimposes every slope; in series mode the sliding window restores the stationarity both need, the gap shrinks to 1.031× vs. 1.006×, and flat-field/SNR remains the safer default because its across-seed spread is the tighter on every regime (±.001 vs. ±.007, ±.003 vs. ±.023 and ±.014 vs. ±.018).

### Calibration, window extent and backend accuracy

Radon velocimetry is sensitive to its calibration: velocity scales linearly with both pixel pitch and frame rate, so an implementation that drops either is off by that ratio. This is the motivation for the synthetic benchmark — it exercises the calibration end-to-end and catches omissions that leave no in-vivo signature. Two details of the sinogram itself matter as much, and both left no in-vivo signature either. First, the window extent: an earlier version of this tool called the Radon transform with scikit-image’s default, which restricts it to the inscribed circle, so every “64-frame” window was in fact its central 32 × 32 block. On the synthetic benchmark the series-mode ratios of Table 4 moved by at most 1.5 percentage points when the whole window replaced the crop, but on real data the whole window is more repeatable: two adjacent 32-px columns of one vessel carry the same blood at the same instant, and on each of the five real curves we examined the median absolute difference between their per-window velocities fell when the whole 64-frame window was used, from 335 µm*/*s to 496 µm*/*s to 302 µm*/*s to 418 µm*/*s (paper_experiments/improvements/e8_ real_repeatability.py). Second, the projector: an earlier GPU backend rotated the window in the opposite sense to scikit-image and summed along the other axis, so the two backends were not evaluating the same angle, and near grazing angles that moved the velocity by tens of percent (one representative column gave 783 vs. 1322 µm*/*s). Because the angle–velocity map is steep near the 85^◦^ bound, a half-degree error from an approximate sinogram moves the velocity by tens of percent, so the sinogram must be computed accurately, not merely quickly; the released GPU projector reproduces the scikit-image sinogram to within 10^−4^ relative, consistent with its single-precision arithmetic (Section 4.7).

### Limitations

Nine remain. First, the patch stage is rigid by construction and assumes a global-shutter imager; genuine intra-frame distortion is corrected only by the B-spline stage, and only at low frame rates. Second, the centroid reference-frame rule (Equation (2)) finds the most central frame, not the best-conditioned one, and can pick an impoverished frame when the eye is mostly closed. Third, the velocimetry needs the operator to trace the vessel center-line by hand, and the sensitivity of the recovered velocity to that tracing — its repeatability within and between operators — was not measured here. Fourth, the per-frame velocities depend on the occlusion (eye-open) mask: closed-eye frames have no nearby valid windows, so the released tool holds the nearest valid velocity (bounded interpolation) and clamps to the velocity-search ceiling (*v*_max_) rather than extrapolating to non-physical values; such frames are also excluded from every reported statistic by the eye-open mask — but a mis-flagged occlusion would corrupt that frame’s velocity. Fifth, the estimator has no defence against a broad intensity band that *moves* across the field. The flat-field step divides each window by a rank-one separable model, which removes a static illumination pattern but not a moving one: inside a 32-px column a sweeping band looks like a gentle spatial ramp whose offset drifts from frame to frame, a broad sloped structure that the angle metric scores exactly as it scores a cell streak, and a band crossing the 2048-px field in half a second reads as roughly 2700 µm*/*s. The effect is global to the image rather than local to a vessel: in XC02, 12 separate instants have all 15 independent vessel columns at the search ceiling at once, which 500 shuffles of each column’s own time course never reproduce (at best they put 53% of columns there together). In the three acquisitions analysed here its ceiling-level form sits at blink boundaries, where the eyelid margin and the tear film sweep across the field; that localisation is a property of these recordings rather than a general rule, and the census below shows the milder form occurring away from blinks in other recordings. Over the window set the tool measured before the any-blank-frame guard (improvements/e15_moving_bands.py), the ceiling rate climbs with how much of the window is blank — 19.5%, 34.7%, 57.1% and 63.0% for windows 0–5%, 5–15%, 15–30% and 30–70% blank — while not one of the 6731 windows with no blank frame reached it and every one of the 471 ceiling windows contained at least one. Those windows are also broad where ordinary ones are not (broad-energy share 0.843 against 0.247), each one tracks the broad component rather than the fine one, and their fine component alone reads a median 445 µm*/*s instead of 2974 µm*/*s. Dropping any window containing a blank frame therefore removes this population entirely, at a cost of 855 of 7586 windows (11%), and that is what the released tool does. A weaker version survives between blinks and is *not*corrected: among the windows the guard keeps, a window’s broad-energy share still correlates with its recovered velocity (Spearman *ρ* = +0.29) and the median velocity rises from 741 µm*/*s to 1128 µm*/*s across quartiles of that share. Part of that is genuine — a truly fast streak is also spatially broad — so it bounds the residual artifact rather than measuring it. How often that form occurs is a property of the recording, not of the subject. A per-frame census of broad structure in the difference video (improvements/e15c_band_census.py) finds bursts of it in all ten recordings of one further cohort subject, at rates that differ across that subject’s own sessions: 0–7 bursts away from blinks per recording on the first visit and 9–19 on the second. The bursts also track eye motion: 86% of the 71 of them that fall away from blinks coincide with a net shift of the registered eye position larger than 50 px, against 23% of random eye-open moments, while how often a recording contains such shifts does not predict its burst rate (Spearman *ρ* = −0.42, *n* = 10). That is the signature expected of a pattern which is not attached to the retina: registration holds the retina still, so the illumination falloff, a tear-film interference fringe or any other structure fixed in the instrument is swept across the registered frame whenever the eye moves, and the frame differencing renders the sweep as a band. We neither detect nor correct the band itself; a per-frame band detector, a guard on windows that span a large registered eye movement, or an angle metric insensitive to low-spatial-frequency structure, is future work. Sixth — and the one that scales every velocity we report — the pixel pitch is a single nominal *p* = 0.67 µm*/*pixel applied to every eye, derived from the instrument’s 5^◦^ field at 2048 samples. The true retinal pitch depends on the individual eye’s axial length, and correcting for it requires a per-eye biometry measurement [45], which is not available for the whole cohort because the biometer was installed part-way through data collection. In healthy adults the axial length typically varies by about ±2 mm around the nominal 24 mm, which scales the retinal pitch — and therefore every reported velocity — by roughly ±8%. This is a systematic per-subject scale factor, not added noise: it does not affect the shape of a velocity waveform, the pulsatility indices, or any comparison within one eye, but it does bound the accuracy of absolute velocities across subjects, and it is one plausible contributor to our means sitting below the published parafoveal values. Per-eye correction is a drop-in change to one calibration constant once biometry is available. Seventh, we have not measured the registration → velocimetry coupling directly: the per-window SD we can quote (3–21 µm*/*s on the constant synthetic) is the estimator’s intrinsic noise floor on an ideal space–time image, a lower bound rather than a registration-calibrated threshold. Eighth, the synthetic benchmark isolates the estimator in two ways that flatter it: the generator adds no background noise (its only noise is the Poisson noise of the cells themselves), and the benchmark feeds the estimator rows of the raw synthetic video rather than the frame-difference stack the pipeline builds from real data, so the postprocessing is never exercised. It is also run on the full 64-px synthetic vessel width; at the 32-px column width used on real data the same estimator reads 1–9% low on these noise-free videos (series-mode ratios 0.989 / 0.908 / 0.994 / 0.961 for constant / linear / pulsatile / constant at 0.67 µm*/*pixel, 20 seeds each). Ninth — and, with the pixel pitch, the other systematic factor on every mean we report — the postprocessor’s 3-D Gaussian (*σ* = 1 frame in time as well as 1 px in space) tilts every streak towards the time axis and so biases the velocity low. Pushing the same synthetic videos through the production postprocessing lowers those ratios to 0.946 / 0.669 / 0.891 / 0.850, an underestimate of 5–33% that depends on the flow regime. Simply removing the temporal component is not the answer on real data: on one 400 Hz acquisition rebuilt without it (322 columns, measured with one whole-window variant of the estimator throughout), the median per-column velocity roughly doubled (755 µm*/*s with the production blur, 1520 µm*/*s with no blur, 1694 µm*/*s with a per-frame 2-D blur) while the agreement between adjacent columns of the same vessel halved (correlation 0.50 → 0.25). The temporal blur is also whitening frame-to-frame noise that the angle metric would otherwise read as near-horizontal streaks, i.e. as very fast flow; we therefore keep it, report the bias, and leave a noise-whitened angle metric as future work. Together with the pixel-pitch factor, this bias is a second reason our means sit below the published parafoveal values.

## 6. Conclusion and future work

We presented an end-to-end pipeline that measures erythrocyte velocity in retinal capillaries from high-speed AO video: an occlusion-aware preprocessor; a GPU patch-based translation stage with RANSAC robust aggregation and automatic reference-frame selection; an optional hierarchical elastix refinement that composes transforms instead of resampling; a frame-difference postprocessor; and a Radon velocity stage implementing the flat-field angle method of Duncan et al. in Python, with CPU and GPU backends that return the same velocities. Both surfaces share one analysis core behind a 3D Slicer extension and a batch tool. On synthetic ground truth the flat-field estimator recovers the mean velocity to within ±4% with a pixel-pitch calibration verified to ±2%; on three 400 Hz acquisitions, taken as a feasibility demonstration, the recovered mean velocities (*V_m_* ≈ 0.95 mm/s) are physiologically plausible (of the same order as published AO values, though below them); registration improves inter-frame consistency across the full 2344-video, 57-subject cohort (SSIM 0.73 → 0.86); and on the 20-acquisition head-to-head the GPU patch-based translation stage aligns the large eye motion that default-pyramid itk-elastix misses, 3.6–38× faster than the best-tuned elastix configuration and with the fewest unaligned frames.

### Future work

Six directions extend the system. *Automated vessel segmentation*, proposing candidate curves from the mean-difference projection, would remove the last manual step and enable whole-field velocity maps. *Cross-instrument validation* against an independent modality (e.g. laser-Doppler flowmetry on a controlled phantom) would establish agreement beyond accuracy against a synthetic signal. A *quantitative head-to-head* against strip-based ophthalmic registration [16, 17] on the same acquisitions would quantify the benefit of whole-frame refinement on the downstream velocity. *Cardiac-cycle timing parameters* — systolic and diastolic landmarks, and the secondary peak reported from AO velocity waveforms by Gu et al. [1] — are a natural next analysis: additional maxima within each cardiac cycle are present in our series and survive the 10 Hz post-filter, so the information is not smoothed away. What is missing is cycle alignment. Gu et al. recorded a finger pulse-transducer signal simultaneously with the video and measured every parameter from a velocity waveform averaged over multiple cardiac cycles; our acquisitions carry no pulse or ECG channel, so this first needs either a reference signal or an agreed procedure for locating cycle boundaries in the velocity trace itself. A *per-frame image-quality score* would generalize the window guard from “is this frame blank” to “is this frame trustworthy”, which is what the moving-band artifact really asks for: the registration stage already fits each frame’s displacement by RANSAC and so already knows how many of its patches voted for the winning shift, a number it currently computes and discards, and a sweeping band should depress it exactly where the velocity spikes. The same stage already records each frame’s displacement, and since most bands away from blinks coincide with a large eye movement (Section 5), a window guard on that displacement is the cheapest partial defence and needs no new measurement. Finally, the temporal component of the postprocessing blur biases every velocity low while also suppressing the frame noise that the angle metric would otherwise misread; a *noise-whitened angle metric* that keeps the second effect without the first would remove a known systematic error from every mean we report.

## Footnotes

1 The registration parameter values quoted in Section 3.3 are those of the current pipeline build; the in-vivo velocity series shown here were produced with an earlier build of the same two-stage registration. Both reduce residual inter-frame motion to the sub-pixel regime the space–time construction requires, so the recovered velocity is insensitive to the difference.

